# Inferring Neuron-level Brain Circuit Connection via Graph Neural Network Amidst Small Established Connections

**DOI:** 10.1101/2023.06.29.547138

**Authors:** Guojia Wan, Minghui Liao, Dong Zhao, Zengmao Wang, Shirui Pan, Bo Du

## Abstract

**Motivation:** Reconstructing neuron-level brain circuit network is a universally recognized formidable task. A significant impediment involves discerning the intricate interconnections among multitudinous neurons in a complex brain network. However, the majority of current methodologies only rely on learning local visual synapse features while neglecting the incorporation of comprehensive global topological connectivity information. In this paper, we consider the perspective of network connectivity and introduce graph neural networks to learn the topological features of brain networks. As a result, we propose Neuronal Circuit Prediction Network (NCPNet), a simple and effective model to jointly learn node structural representation and neighborhood representation, constructing neuronal connection pair feature for inferring neuron-level connections in a brain circuit network.

**Results:** We use a small number of connections randomly selected from a single brain circuit network as training data, expecting NCPNet to extrapolate known connections to unseen instances. We evaluated our model on *Drosophila* connectome and *C. elegans* worm connectome. The numerical results demonstrate that our model achieves a prediction accuracy of 91.88% for neuronal connections in the *Drosophila* connectome when utilizing only 5% of known connections. Similarly, under the condition of 5% known connections in *C. elegans*, our model achieves an accuracy of 93.79%. Additional qualitative analysis conducted on the learned representation vectors of Kenyon cells indicates that NCPNet successfully acquires meaningful features that enable the discrimination of neuronal sub-types. Our project is available at https://github.com/mxz12119/NCPNet.

## Introduction

Understanding how the interconnections of neurons account for brain functions has been a great pursuit over the past decades [Eichler et al., 2017, Lichtman and Denk, 2011, Liu and Qian, 2022, Dey et al., 2022, Wang et al., 2022]. A brain circuit network refers to a physical map of a brain’s structural connections, commonly referred to as a connectome. This connectome naturally lends itself to a graph-theoretic representation, wherein neurons are depicted as nodes and their interconnections are depicted as edges. One critical barrier that has hindered a comprehensive understanding of the human brain circuit network is the limited scalability of neuronal connection reconstruction techniques.

A typical synapse-level connectome reconstruction pipeline [Takemura et al., 2015] includes Electron Microscopy (EM) imaging, neuron segmentation[Sheridan et al., 2023], 3D reconstruction [Januszewski et al., 2018] and synapse prediction [Buhmann et al., 2021]. Although automatic approaches based on computer vision (CV) greatly speed up tracing neurons, but reconstructing millions of synapse connections has remained a laborious hand-craft task [Dorkenwald et al., 2022]. Existing approaches concentrate on extracting local synaptic features such as T-bars and synaptic clefts using Convolutional Neural Networks (CNNs) [Buhmann et al., 2021]. However, CNNs ignore the global topological information over a graph-like connectome for predicting neuronal connections. The graph-like interactions that intricately shape the functioning [Rubinov and Sporns, 2010] of a nervous system inherently dictate neuronal connections. Thus, there is an ongoing requirement for an efficient methodology capable of capturing and representing these essential topological features.

In the recent years, Graph Machine Learning (GML) has seen a great surge of attention with Graph Neural Networks (GNNs) being successfully developed due to their powerful expressiveness for analyzing graph-structured data, benefiting a myriad of applications including social network [Andreassen, 2015], drug discovery [Satorras et al., 2021], etc. GNNs learn node representations for a graph by following a recursive message passing scheme, similar to the behaviors that neurons or brain function regions intercommunicate. Instead of modeling neuron-level networks, recent advances mainly concentrate on region-level human brain networks, of which brain atlas is constructed from a set of regions of interest (ROI) as nodes by functional Magnetic Resonance Imaging (fMRI) techniques [Kan et al., 2022, Li et al., 2021, Kim et al., 2021]. To date, the investigations of learning neuron-level connections in term of graph machine learning are still inadequate. Recently, main stream approaches of neuron synaptic connection prediction are based on visual synaptic features or neuron segmentation [Buhmann et al., 2021, Turner et al., 2020]. In the earlier year, [Kreshuk et al., 2015] propose a probabilistic graphical model to predict potential connections. As a non-learnable model, however, the approach could be hard to generalize effectively when confronted with more complex situations.

A prospective strategy might involve the development of graph-based machine learning models intended for learning the structure feature of a brain circuit network. We present our idea in Figure 1 that neuronal connection prediction can be feasibly modeled as a link prediction task. As a classic task in data mining [Martínez et al., 2016], link prediction is employed to predict the likelihood of a connection existing between two nodes within a given graph.

**Fig. 1.**
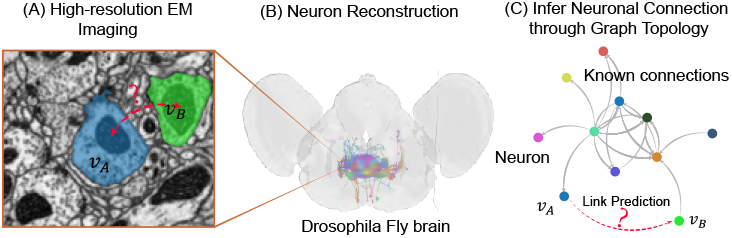
The schematic illustration of inferring potential neuronal connections between *v*_*A*_ and *v*_*B*_ through graph topology.

link prediction is to infer whether two nodes in a graph are likely to have a connection. GNNs have been proceeded a successful way with achieving high accuracy in link prediction by jointly learning graph structure information. However, it is non-trivial to apply such frameworks on brain circuit networks due to domain differences. In specific, a brain circuit network contains rich motifs that appear in characteristic frequencies [Sporns et al., 2004]. A motif acted as a simple circuit in a brain network is a subgraph pattern repeated and reused many times for performing a specific functional role [Sporns et al., 2004]. Some recent works have theoretically proven that subgraph isomerism issue [Xu et al., 2018] reduce the representational capacity of the aggregation-based GNNs [Xu et al., 2018]. In particular, it becomes more worse for link prediction [Srinivasan and Ribeiro, 2020].

In this paper, we presents the application of Graph Neural Networks (GNNs) to neuron-level connectomes, with the objective of predicting neuronal connections in an end-to-end manner. For this, we propose Neuronal Circuit Prediction Network (NCPNet) to jointly learn node representation and neighbor representation over brain circuit networks. Our model contains two components. One is Node Structural Information Encoder (NSIE), which leverages Graph Convolutional Network to learn latent node representations. Another is Neighborhood Encoder (NBE), which learns neighbor-shared structure information to distinguish neighboring nodes. Then we fuse two representations by a max-pooling layer and calculate connection similarity by a fully connected layer. For optimizing the model, we leverage negative sampling to minimize the cross entropy loss. To evaluate our model, we perform link prediction on the *C. elegans* worm connectome [White et al., 1986, Cook et al., 2019] and the *Drosophila* hemibrain connectome [Scheffer et al., 2020]. Experimental results show that our model achieves excellent performance for predicting unseen connections, which achieves 98.15% AUC on the *Drosophila* connectome and 98.59% on the *C. elegans* connectome. To our surprise, we find that even 5% training connections can achieve over 90% accuracy.

Our contributions are summarized as following:

- We propose NCPNet, an end-to-end model for capturing meaningful topological feature for brain circuit networks. This represents a new investigation into the application of Graph Neural Networks in neuron-level connectomes to predict neuronal connections.
- NCPNet exhibits remarkable predictive prowess for neuronal connections, even when trained only on a small fraction of the total connections.
- By utilizing the learned topological features, NCPNet is capable of distinguishing not only the functional types but also the sub-types within a group of neurons.

## Preliminaries and Task Definition

### Brain Circuit Network

In this paper, we only consider neuron-level connections. Let a brain circuit network be a directed graph *G* = (*𝒱, ε*), where *𝒱* = *{v*_1_, *v*_2_, *· · ·, v*_*N*_ *}* is a set of nodes, *ε ⊆ 𝒱 × 𝒱* is a set of edges, and the total number of nodes is *N*. *v ∈ 𝒱* denotes a neuron. (*v*_*i*_, *v*_*j*_) *∈ ε* denotes a connection from the pre-synaptic neuron *v*_*i*_ to the post-synaptic neuron *v*_*j*_. The adjacency matrix of a brain circuit network is **A** *∈* ℝ^*N×N*^, where *A*_*ij*_ = 1 if (*v*_*i*_, *v*_*j*_) *∈ ε* and 0 otherwise. Let *P* (*Y* |(*v*_*i*_, *v*_*j*_)) denotes the probability that *v*_*i*_ connects to *v*_*j*_. Each node *v*_*i*_ has its own feature vector *x*_*i*_ *∈* ℝ^*d*^, with **X** *∈* ℝ^*N×d*^ denoting the collection of all node feature vectors in a matrix form. Let Γ_*k*_(*v*_*i*_) be the *k*-hop neighbors of *v*_*i*_. The degree matrix **D** *∈* ℝ^*N×N*^ is a diagonal matrix defined by *D*_*ii*_ = |Γ_1_(*v*_*i*_)|.

### Neuronal Connection Prediction

In this paper, we use (*v*_*i*_, *v*_*j*_) to denote a connection from *v*_*i*_ to *v*_*j*_. (*v*, ?) means ‘which neurons are post-synaptic for *v* ?’. Similarly, (?, *v*) means ‘which neurons are pre-synaptic for *v* ?’. Given a brain circuit network *G, ε*_+_ *⊂ ε* is the known connections and **X** is the node feature matrix. Let *ε*_+_ and *X* be training data, neuronal connection prediction aims to predict *P* (*Y* |(*v*_*i*_, *v*_*j*_), (*ε*_+_, *X*), *θ*) = *f*_*ε*_ (*v*_*i*_, *v*_*j*_) by a computational model *f* under the parameter *θ*.

## Method

We now introduce NCPNet for neuronal connection prediction as illustrated in Figure 2. For the input brain circuit network *G* and the node feature *X*, our model can predict a given connection (*v*_*i*_, *v*_*j*_) *∈ 𝒱 × 𝒱*. We take a positive connection (*v*_*A*_, *v*_*B*_) as an example displayed in Figure 2. It consists of two components:(1) Node Structural Information Encoder (NSIE) and (2) Neighborhood Encoder (NBE).

**Fig. 2.**
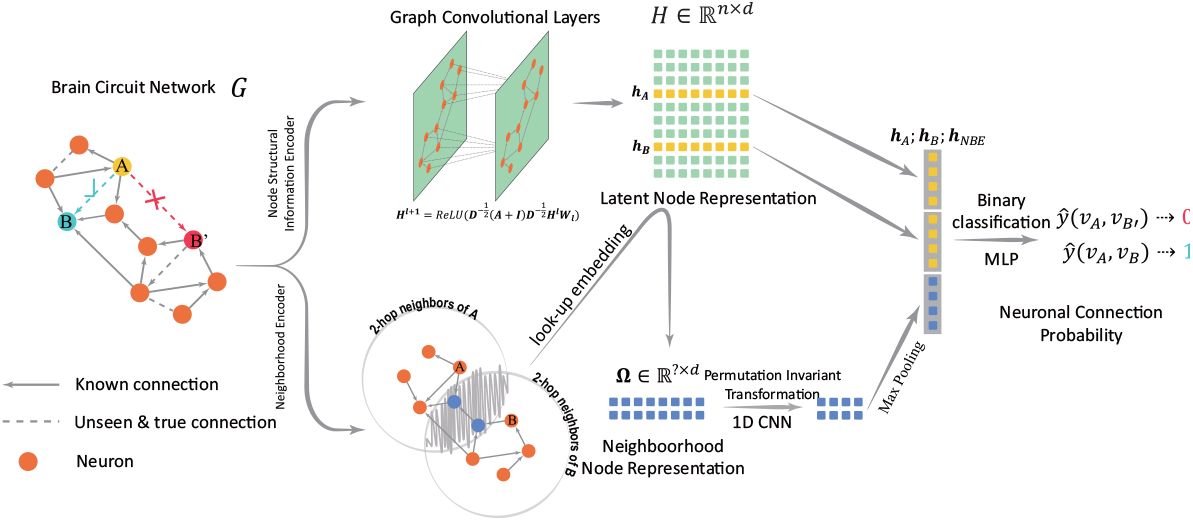
The architecture of NCPNet for link prediction. Our model consists of two modules: 1) the top workflow depicts Node Structural Information Encoder with the input of the adjacency matrix **A** and the feature matrix **X**, and output the latent node representations **H**; 2) the bottom depicts Neighborhood Encoder to learn pair node neighbor representation to allow link prediction distinguishable, and output neighborhood representations. Finally, the two kinds of representations are fed into the max-pooling layer and a MLP layer to score neuronal connection probability.

### Node Structural Information Encoder

NSIE can optional receive the input of node attribute information, such as neuron type information, neuron location. In this paper, we specifically focus on neuron type information provided by [Scheffer et al., 2020]. A detailed illustration of neuron type information can be seen in Part 1 of the supplementary materials (SI). We transform such type information into an one-hot feature matrix **X** *∈* ℝ^*N×m*^. Next, we use a linear layer to project **X** into a continuous feature matrix **H**^(0)^ = **X** *·* **T**, where 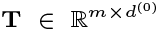 denotes type embedding, *m* is the number of neuron types and *d*^(0)^ is the dimension of the initial feature matrix. If type information is not included, **H**^(0)^ is initialized using Glorot initialization. Subsequently, we feed the adjacency matrix **A** and the feature representation **H**^(0)^ into GCN layers for learning latent features:

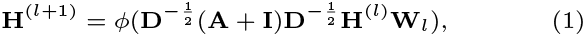

where 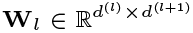 is a trainable projection matrix of *l*-th layer. **I** *∈* ℝ^*N×N*^ is an identity matrix. *φ* is ReLU function. We only stack 2 layers in the architecture because more layers lead to over-smoothing of node representations [Chen et al., 2020].

Following the iterative learning process of the GCN layers, the node representation **h**_*A*_ and **h**_*B*_ can be indexed from the final latent feature matrix for predicting the connection (*v*_*A*_, *v*_*B*_).

### Neighborhood Encoder

To improve the expressiveness, we propose Neighborhood Encoder (NBE) to capture neighborhood overlap information. Neighborhood information has proven important for link prediction [Yun et al., 2021]. The addition of neighborhood information makes edge predictions distinguishable because neighborhood information is determined by a node pair rather than a single node.

We define the neighborhood information of a neuronal connection as

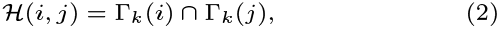

where *ℋ* (*i, j*) is the set of *k*-th hop neighbors of node *v*_*i*_ and *v*_*j*_. We set *k* as 2 for avoiding including too much unrelated nodes. Then the neighbor node representation is obtained by looking-up embeddings from the last output **H** of NSIE:

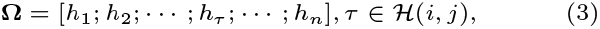

where ‘;” denotes vector concatenation, thus **Ω** *∈* ℝ^| *ℋ* (*i,j*)|*×d*^.

Next, we leverage 1D CNN with a stride of 1 to learn neighborhood features. Such operation is suitable for learning elements in a set due to its permutation invariance:

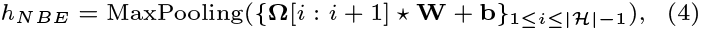

where 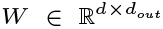 is a 1D CNN kernel, and 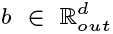 is bias. *\** denotes 1D convolution operator. MaxPooling(*·*) denotes element-wise max pooling, reducing neighborhood features into 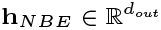. Importantly, when stride=1, the above function is permutation invariant to any transposition of *h*_*i*_ and *h*_*j*_ in Ω _*ℋ*_.

Finally, we concatenate **h**_*A*_, **h**_*B*_ and **h**_*NBE*_ as a joint representation of (*v*_*A*_, *v*_*B*_). For learning the neuronal connection asymmetry, we use a 2-layer fully connected layer as an asymmetric predictor to calculate scores, which means *P* (*Y* |(*v*_*A*_, *v*_*B*_)) ≠ *P* (*Y* |(*v*_*B*_, *v*_*A*_)). The predictor is written by

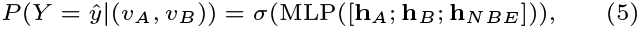

where; denotes concatenation operator. *σ* is a sigmoid function.

### Negative Sampling Training

In practice, the training data *ε*_+_ *⊂ ε* only contains positive connections. But the model need negative connections for building the capability of distinguishing positive and negative connections. A conventional strategy is randomly sampling negative edges to augment the training data. In subsequent, the training procedure can be expressed by such loss function

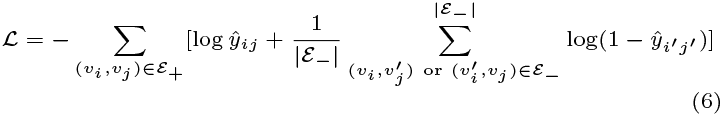

where *ε*_*−*_ denotes the negative samples. 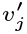 or 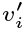 are randomly chosen from *𝒱* to construct negative edges. | *ε*_*−*_| denotes the number of negative samples.

## Experiments and Results

### Quantitative Analysis on Neuronal Connection Prediction

#### Brain circuit network data preparing

In this work, we use two connection matrices released by *Drosophila* [Scheffer et al., 2020] and *C. elegans* [Cook et al., 2019]. The brief introductions of the two connectome are listed as following.

- *Drosophila* hemibrain is now the largest synaptic-level connectome, which covers half part of the central fly brain [Scheffer et al., 2020]. It was reconstructed from over 20 TB High-resolution EM images. The original data is available at ^1^. We pre-processed ‘hemibrain version 1.2.1’ connectome for adapting graph machine learning tasks. The pre-processed dataset is released at our code repository ^2^.
- The *C. elegans* connectome is reconstructed from both sexes of nematodes [Cook et al., 2019], of which raw data is available at ^3^.

Each of them can be depicted as a directed graph. The statistics of them can be seen in Table 1. The *C. elegans* connectome belonging to a male nematode has 598 nodes with 7813 edges deriving from chemical connection or electrical connection. The *Drosophila* connectome has 21793 neurons with about 355.04 k edges deriving from synaptic connections. As shown in Table 1, both *Drosophila* and *C. elegans* are sparse graphs with respectively 0.75% and 2.19% of all possible edges. Both they have only one connected component (Figure S1,S4). As Figure S3, S6 of SI show, they both approximate truncated power-law distributions, indicating that a neuron tend to build connections to others beyond a specific threshold. This property differs from social networks and citation networks [Kossinets and Watts, 2006]. Besides, the two brain circuit networks *Drosophila* both present high clustering and low average path length, implying a small-worldness that most nodes can be reached from every other node by a small number of hops.

**Table 1.** The statistics of the brain circuit networks used in this paper, where ‘DG’ denotes average degree, ‘SP’ denotes average shortest path length, ‘CC’ denotes average clustering coefficient, and ‘CP’ denotes characteristic path length.

| Datasets | $\vert V\vert $ | $\vert \mathcal{E}\vert $ | Type | Density | DG | SP | CC |
| --- | --- | --- | --- | --- | --- | --- | --- |
| Drosophila | 21793 | 355.04 k | 5554 | 0.0075 | 326.6 | 2.7305 | 0.2382 |
| C.elegans | 598 | 7813 | 19 | 0.0219 | 26.13 | 3.7029 | 0.2557 |

#### Experimental settings

For simplifying models, we only regard the two brain circuit networks as directed graphs without modeling synapse strength feature. We employed a methodology of varying the size of the training set to assess the model’s performance under different known connections. Specifically, we partition the total edges in *E*_+_ of each brain circuit network into various segments: x% is allocated for training models, 10% is dedicated to validation and optimization of hyper-parameters, while the remaining (90-x)% is reserved for testing. Within the test set, we adopt a similar approach to the negative sampling described in Section 4, generating an equal number of negative connections to match the positive ones.

Neuronal connection prediction can be modeled as a link prediction task in graph machine learning. It can be regarded as binary classification that aims to distinguish positive connections and negative connections by a classification threshold. The best classification threshold of every model is determined by maximizing accuracy performance. Then we use AUC, F1, Accuracy, and Recall as quantitative metrics to evaluate our model.

We compare our model with 5 representative link prediction methods: Common Neighbors (CN), Adamic Adar (AA), Node2Vec [Grover and Leskovec, 2016], GCN [Kipf and Welling, 2017], SEAL [Zhang et al., 2021]. Among them, CN and AA are heuristic Methods, and Node2Vec is a representative graph embedding method. GCN is the most famous graph neural networks. We conducted 10 independent runs and calculated their mean and the standard deviation.

We leverage Pytorch Geometric to implement NCPNet and other GNN methods including GCN [Kipf and Welling, 2017] and MLP. We use the code of SEAL from the referenced paper [Zhang et al., 2021]. For graph embedding methods and Node2Vec [Grover and Leskovec, 2016], we obatain node embeddings by GraphVite ^4^ and use dot product to calculate connection similarity. For heuristic methods, Common Neighbors (CN), Adamic Adar (AA), we use Networkx ^5^ to predict connection scores, which is a popular open-source graph toolkit.

All experiments are conducted on GPU as NVIDIA GeForce RTX 3080Ti (12 GB) and NVIDIA TITAN Xp (12 GB), CPU as Intel(R) Xeon(R) CPU E5-2620 2.10GHz. For fair comparison, we reported results of all baselines and NCPNet as the mean and the standard deviation of performances from 10 independent runs.

For hyperparameter settings, we grid-searched all hyper-parameters and the optimal settings are shown in Table S1 of SI.

#### Neuronal Connection Prediction Results

The connection prediction results are displayed in Table 2 and Figure 3A. We can observe from Table 2 that NCPNet consistently outperforms other baselines on two datasets with varying training edges. In specific, comparing CN and AA to the other data-driven methods, CN and AA exhibit poor performance when trained on a limited number of edges (*<* 10%), but comparable performance over *>* 30% training edges, indicating that such heuristic methods do not capture complex topological feature well. Importantly, our model presents significant performance using low training data. To our surprise, even using 1% training connections (about 35.5 k connections of *Drosophila*), NCPNet achieves 94.43% AUC, 88.59% accuracy, 88.82% F1, and 90.80% recall. As shown in Figure 3A, GNN models (NCPNet and GCN) show a sharply increase when training data is less than 5%(Figure 3(I, II), *Drosophila*) or 10% (Figure 3(IV, V), *C. elegans*), implying the efficacy that GNNs can learn the structure information of brain circuit networks. However, the performance curves of GCN become saturated after 10% seen in Figure 3A(I, II,IV, V). This suggests that conventional GNNs reach a limit in their ability to capture the expressive power required for learning brain circuit networks. Some theoretical studies on GNNs reveal that an abundance of motifs with repetitive occurrences in a graph can diminish the expressive capacity of GNN models [Xu et al., 2018], thereby resulting in deceptive outcomes during link prediction [Srinivasan and Ribeiro, 2020]. By incorporating Neighborhood Encoder, NCPNet can acquire the ability to discern isomorphic nodes through the learning of overlapped neighborhoods. Consequently, NCPNet demonstrates a substantial enhancement over the performance of GCN.

**Table 2.** Neuronal connection prediction results(AUC). **Bold** indicates the best performance in a column. Each number is the average performance for 10 random runs of the experiments. We also report data storage size used in the training process.

| Method | *Drosophila* |  |  |  | *C. elegans* |  |  |  |  |
| --- | --- | --- | --- | --- | --- | --- | --- | --- | --- |
|  | 1% | 5% | 10% | 50% | 1% | 5% | 10% | 50% |  |
| Data Size | 0.35 MB | 1.44 MB | 2.79 MB | 13.62 MB | 12.94 KB | 5.39 KB | 8.44 KB | 32.85 KB |  |
| CN | 50.33±0.15 | 62.34±0.59 | 79.67±0.14 | 94.58±0.11 | 50.47±1.85 | 52.67±0.45 | 60.41 ±1.21 | 89.16±0.73 | AUC |
| AA | 50.92±0.23 | 66.20 ±0.72 | 80.21±0.26 | 94.77±0.06 | 50.21±0.84 | 54.39±0.38 | 60.43±0.97 | 89.66±0.24 |  |
| Node2Vec | 72.71±0.14 | 84.30 ±0.11 | 89.13±0.08 | 93.15±0.09 | 52.96 ±3.42 | 53.83 ±0.69 | 56.52±0.28 | 75.95 ±0.18 |  |
| MLP | 88.47±2.57 | 91.56±0.84 | 94.25±0.56 | 94.89±0.16 | 79.43 ±3.27 | 82.83±1.26 | 84.62±0.57 | 86.29 ±0.23 |  |
| GCN | 89.45±1.24 | 91.56±0.12 | 94.25±0.10 | 95.02±0.07 | 76.01±1.34 | 90.47±0.68 | 91.46 ±0.52 | 97.25 ±0.37 |  |
| SEAL | 87.64±0.26 | 92.18±0.23 | 95.94±0.17 | 96.14±0.15 | 80.56±3.27 | 84.37±0.94 | 88.97±0.68 | 94.34±0.53 |  |
| NCPNet | **94.43±0.08** | **96.50±0.06** | **97.14 ±0.07** | **98.15 ±0.04** | **86.42 ±1.73** | **96.92±0.41** | **97.76±0.53** | **98.59±0.44** |  |
| CN | 50.11±2.14 | 50.83±0.49 | 56.22±0.17 | 88.46±0.11 | 50.00±1.25 | 50.16±0.56 | 52.12±0.41 | 82.27±0.27 | Accuracy |
| AA | 51.03±1.23 | 54.58±0.41 | 66.21±0.21 | 89.34±0.08 | 50.60±0.90 | 52.59±0.41 | 60.29±1.20 | 86.12±0.43 |  |
| Node2Vec | 68.83±0.29 | 76.74±0.41 | 81.10±0.11 | 86.04±0.08 | 52.83±1.77 | 55.71±1.10 | 57.92±0.96 | 72.27±0.26 |  |
| MLP | 80.81±1.85 | 83.94±0.76 | 87.42±0.58 | 88.58±0.12 | 79.34±2.17 | 78.28±0.74 | 83.26±0.62 | 78.06±0.14 |  |
| GCN | 83.70±0.20 | 90.92±0.11 | 90.97±0.17 | 90.99±0.14 | 80.30±1.59 | 83.62±1.36 | 86.68±0.20 | 92.88±0.13 |  |
| SEAL | 84.28±0.24 | 88.53±0.13 | 91.54±0.11 | 91.87±0.12 | 70.42±1.21 | 77.10±0.72 | 83.19±0.69 | 89.02±0.16 |  |
| NCPNet | **88.59±0.18** | **91.88±0.12** | **92.57±0.10** | **93.51±0.07** | **83.10±1.74** | **93.79±1.15** | **96.36±0.80** | **95.51±0.15** |  |
| CN | 0.02±0.31 | 3.71±0.38 | 23.72±0.15 | 89.31±0.09 | 0.00±0.00 | 0.67±0.19 | 8.15±2.20 | 79.59±0.69 | F1 |
| AA | 3.58±0.39 | 11.69±0.24 | 55.28±0.09 | 90.37±0.05 | 2.47±0.98 | 10.31±1.64 | 37.01±1.45 | 85.77±1.65 |  |
| Node2Vec | 67.36±0.43 | 77.22±0.32 | 81.50±0.05 | 86.33±0.04 | 46.88±5.53 | 42.72±3.19 | 50.32±2.03 | 68.78±1.07 |  |
| MLP | 81.05±0.43 | 83.87±0.32 | 87.49±0.16 | 88.60±0.04 | 60.80±0.87 | 63.86±0.80 | 66.01±0.10 | 65.71±0.11 |  |
| GCN | 83.90±0.13 | 91.10±0.09 | 91.21±0.11 | 91.22±0.07 | 61.66±1.18 | 72.48±0.61 | 75.39±0.14 | 86.90±0.17 |  |
| SEAL | 84.15±0.13 | 90.02±0.13 | 91.77±0.07 | 91.91±0.09 | 61.32±0.93 | 66.22±0.65 | 70.67±0.28 | 80.27±0.25 |  |
| NCPNet | **88.82±0.07** | **92.08±0.11** | **92.74±0.06** | **93.76±0.07** | **80.54±1.38** | **88.71±0.84** | **93.02±0.34** | **92.80±0.19** |  |
| CN | 0.05±0.24 | 2.01±0.15 | 13.60±0.11 | 88.03±0.07 | 0.00±0.00 | 0.32±0.13 | 4.32±1.25 | 71.17±2.35 | Recall |
| AA | 1.95±0.26 | 6.31±0.17 | 39.73±0.15 | 92.06±0.09 | 1.22±0.57 | 5.65±1.47 | 22.34±2.17 | 85.00±1.54 |  |
| Node2Vec | 64.45±0.24 | 78.59±0.22 | 83.20±0.12 | 88.20±0.06 | 38.44±5.51 | 36.42±4.27 | 42.37±3.41 | 61.52±2.10 |  |
| MLP | 80.34±1.23 | 83.99±0.55 | 88.11±0.35 | 88.62±0.05 | 62.68±1.34 | 74.36±0.85 | 64.96±1.64 | 84.33±1.12 |  |
| GCN | 85.03±0.19 | 92.88±0.13 | 93.55±0.19 | 93.56±0.05 | 76.00±1.56 | 87.75±1.13 | 81.77±1.29 | 94.87±0.57 |  |
| SEAL | 86.05±0.29 | 90.35±0.14 | 93.87±0.10 | 94.31±0.07 | 92.31±1.47 | 89.17±1.24 | 81.20±0.57 | 89.46±0.33 |  |
| NCPNet | **90.80±0.22** | **94.16±0.17** | **94.89±0.16** | **95.81±0.05** | **96.72±3.42** | **97.44±1.230** | **97.64±0.65** | **98.29±0.35** |  |

**Fig. 3.**
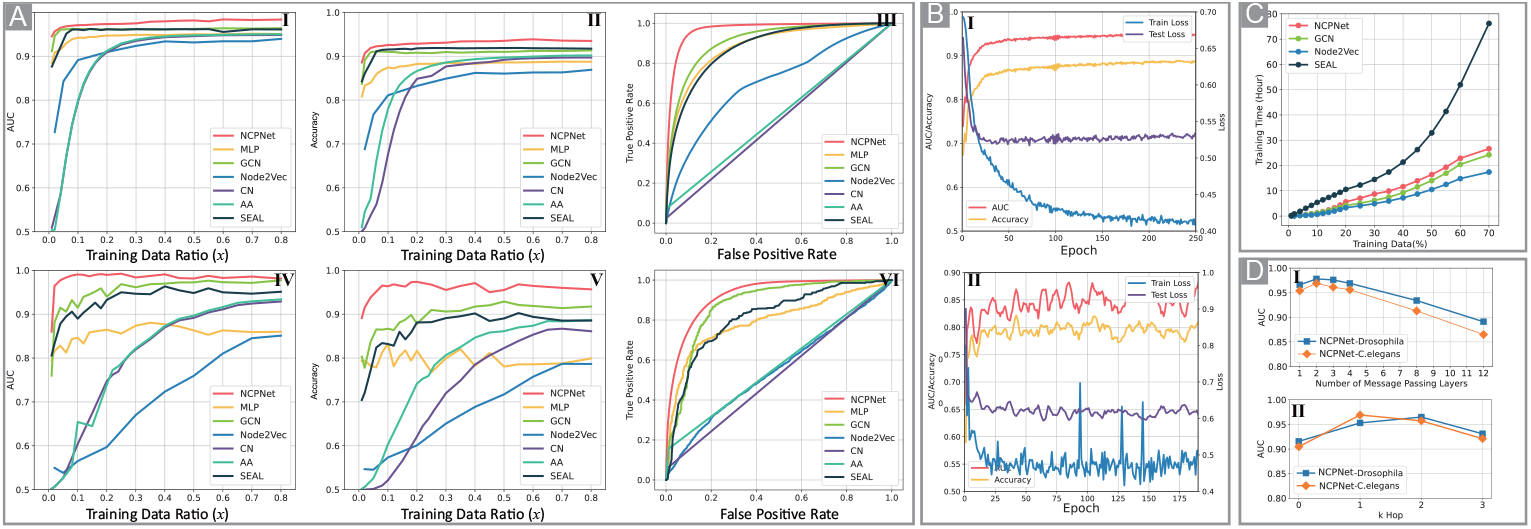
A: Neuronal connection prediction results versus training data ratio. A(I,II,III): The AUC, accuracy, and receiver operating characteristic (ROC) curves of the *Drosophila* connectome. A(IV,V, VI): The AUC, accuracy, and ROC curves of the *C. elegans* conectome. B(I): The training curves of 1% *Drosophila* connectome. B(II): The training curves of 1% *C. elegans* conectome. C: The running time of models on *Drosophila* connectome. D(I): The effect of message passing layers. D(II): The effect of hops for choosing neighbor nodes.

#### Ablation Study

To identify the efficacy of various components within NCPNet, we conducted ablation experiments on the aforementioned model under consistent settings on the 5% *Drosophila* dataset. ‘Type’ refers to the incorporation of neuron type information within the model. As shown in Table 3, all the components are indispensable and contribute to performance improvement. Specifically, we can observe the importance that ‘NSIE’*>*‘NBE’*>*‘Type’.

**Table 3.**
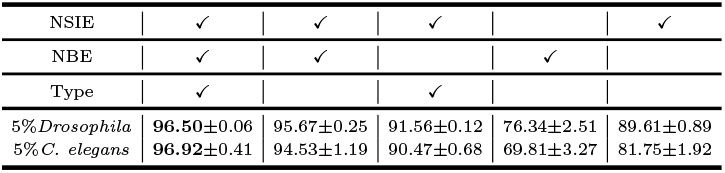
Ablation Study.

#### Model Analysis

##### Training and convergence

We use the same settings from Table 2 and plot the training curves using 1% *Drosophila*/*C. elegans* as shown in Figure 3B(I,II). We can observe that NCPNet converges fast on both two brain networks. The curves of *C. elegans* present obvious fluctuation due to the small amount of training samples on 1% *C. elegans*.

##### Computational complexity and running time

NSIE takes *O*(*N* |*E*|*d* + *Nd*^2^) computational time to compute node representations, which has been well-studied in [Chiang et al., 2019]. *d* denotes the dimension of node representations. Neighborhood Encoder uses the adjacency matrix *A ∈* ℝ^*N×N*^ for locating neighborhood nodes, of which time complexity is *O*(*N* |*E*|_*k*_*d*), where |*E*|_*k*_ is the number of connected edges within *k*-hop. To address this high complexity problem, we convert *A* to a sparse matrix form and thus the complexity reduces to *O*(|*E*|_*k*_*d*). More importantly, we can pre-store *k*-hop neighbor nodes, therefore there is no additional cost to search for neighbor nodes during training and inference.

We also report the practical training time as shown in Figure 3C on *Drosophila*. We only record the training time at the 400-th epoch. Consistent with the above theoretical analysis, similar to GCN, NCPNet shows a linear growth with the number of training edges, indicating that introducing NBE does not increase too much cost for training.

##### Key Hyper-Parameter Analysis

NCPNet involves a number of hyper-parameters. We examine how the different choices of parameters affect the performance of NCPNet on *Drosophila* connetome. As shown in Figure 3D(I), the optimal number of message passing layers is 2. The optimal hop number is 1 for *C*.*elegans* and 2 for *Drosophila*.

### NCPNet Can Learn Meaningful Graph-topological Feature

We reduce the learned representation vectors of every neuron from NCPNet into 2D space by tSNE [Van der Maaten and Hinton, 2008] and plot them in Figure 4 and Figure 5. We further annotate neurons with their types in color. From Figure 4 and 5, we can see that the major portion of representations can form various clusters. Neurons with similar types trend to be close to each other in the latent representation space. This indicates that our model can successfully learn meaningful structural features of brain circuit networks.

**Fig. 4.**
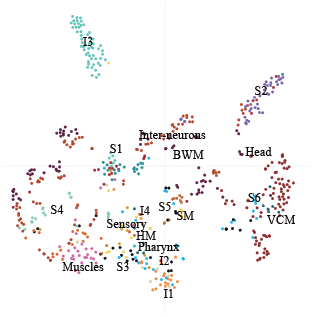
The tSNE visualization of *C. elagans*.

**Fig. 5.**
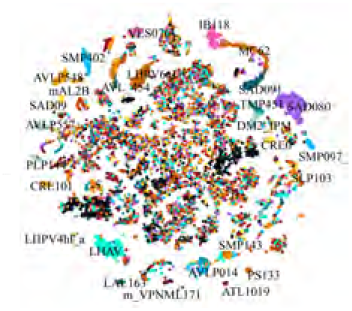
The tSNE visualization of *Drosophila*. We omit partial labels for avoiding over-covering.

### Prediction Score Analysis

The conventional pipeline to identify neuronal connections depends on pre- and post-synaptic features (such as T-bar and synaptic cleft) [Buhmann et al., 2021], which uses image-based neural models to capture these visual features. Rather than using visual features, topology is a prominent and heritable feature of brains. In this section, we apply our model on a typical neuron (ID 509410587) ^6^, which is a protocerebral bridge-ellipsoid body-noduli neuron [Scheffer et al., 2020] shown in Figure 6A.

**Fig. 6.**
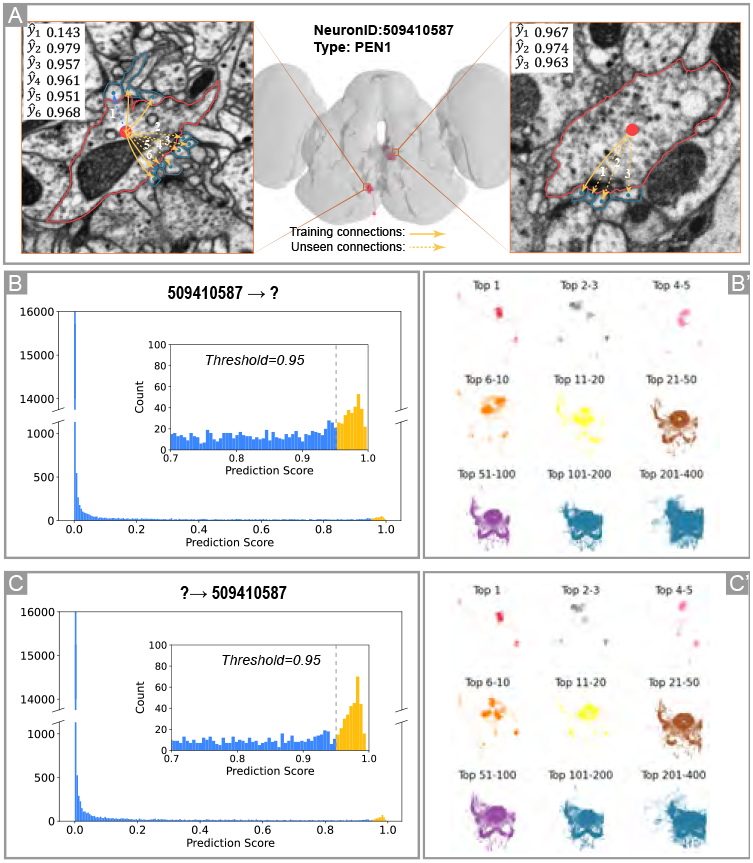
NCPNet prediction scores on *Drosophila* connectome using 5% connections. A: The EM image of Neuron 509410587 in the hemibrain database [Plaza et al., 2022]. We selected cross-sections of EM images of two regions separately to show the local connectivity from the neuron to other neurons. B: The raw score distribution of predicting (*v*_509410587_, ?), in which 509410587 is the pre-synaptic neuron. B’: The 9 groups of the top 400 hits of B. C: The raw score distribution of predicting (?, *v*_509410587_), in which *v*_509410587_ is the post-synaptic neuron. C’: The 9 groups of the top 400 hits of C.

We trained a NCPNet model with 5% known connections, and visualize its predicting confidences on this neuron. As the 3D visualization image (Figure 6A, middle) shows, the neuron mainly consists of two dense dendrite regions across long distance. We selected cross-sections of EM images of two regions separately to show the local connectivity from the neuron to other neurons. Let (*v*_*i*_, *v*_*j*_) be a connection from *v*_*i*_ to *v*_*j*_. (*v*, ?) means ‘which neurons are post-synaptic for *v*’. Our model predicts (*v*_509410587_, ?) to identify the post-synaptic connections with all other neurons. We list the predicting confidences of the connections around these two regions in Figure 6A. It can be observed that our model can precisely predict whether two neurons are synaptically connected. Furthermore, by comparing with two sites shown in Figure 6A, our model presents global predictive capability. This capability is lacking in current methods [Buhmann et al., 2021] that rely solely on local visual synapse features.

Additionally, the distribution of scores is shown in Figure 6B and C. The corresponding top 400 neurons are presented in Figure 6 B’ and C’. The score histograms (Figure 6 B and C) show that only a minority of hits are above the threshold 0.95. We divided the top 400 neurons with 9 groups. Only the highest scoring hits (group I to III) appear to be of exactly the types of pre-synaptic neurons or post-synaptic neurons. Furthermore, we can observe that the prediction results of (*v*_509410587_, ?) and (?, *v*_509410587_) are different, indicating that the predictions are asymmetric just like the directed connection of pre- and post-neurons. In other words, our model can predict pre- or post-synaptic connection separately. More cases can be found in Figure S9-S12 of SI or our code repository.

Existing image-based learning models cost large computation resource to find synaptic connection patterns in 3d volumetric images and suffer from sparsity problem. Our model is lightweight and effective, only depending on a little topology information, and achieves significant performance to measure neuronal connection.

### NCPNet Can Distinguish Kenyon Cell Sub-Types

Next, we investigate the potential of our model to facilitate the clustering of neurons, thereby unveiling functional classes. Specifically, our focus lies on Kenyon cells (KCs), which serve as intrinsic neurons within the mushroom body (Figure 7A(1)), and play a pivotal role in the processes of memory formation and retrieval [Kahsai and Zars, 2011]. KCs form the medial lobe, consisting of the *γ, β*^*I*^, and *β* lobes, and the vertical lobe, consisting of the *α* and *α*^*I*^ lobes (Figure 7A(2)).

**Fig. 7.**
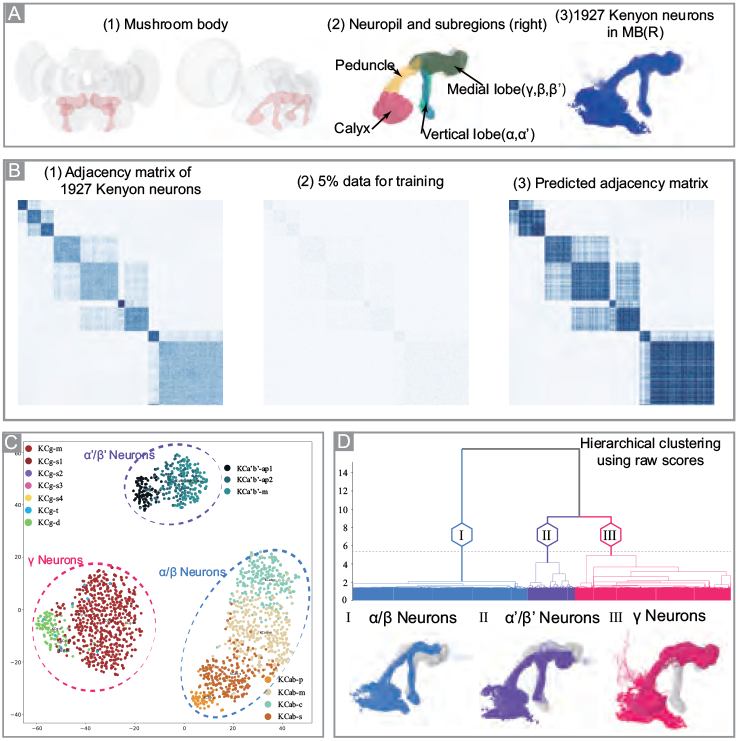
Investigation on distinguishing functional type and sub-types of Kenyon Cells. A(1): Mushroom body in a fly (*Drosophila*) brain. B(2): The right half of the mushroom boy and its sub-regions. A(3): 1927 Kenyon cells in the right mushroom body. B(1): The adjacency matrix of 1927 Kenyon cells. B(2) The 5% connections of B(1). B(3): NCPNet predicts the scores of each Kenyon cells against all others. Let scores *>* 0.95 be true and then a predicted adjacency matrix is obtained. C: The tSNE visualization of the embeddings of KCs. (purple: *α* ^*′*^*/β* ^*′*^ neurons; red: *γ* neurons; blue: *α/β* neurons). D: Hierarchical clustering results and its correlate neurons shown below the dendrogram.

Figure 7A(3) shows the right half of mushroom body containing 1927 KCs in the fly hemibrain, representing 8.8% of the *Drosophila* dataset. The adjacency matrix of these neurons is shown in Figure 7B(1). In our study, we trained our model using a mere 5% of the hemibrain connections, without incorporating any neuron type information. Those scores exceeding the threshold of 0.95 were considered true, leading to the formation of a predicted adjacency matrix seen in Figure 7B(3). Notably, our model exhibits the ability to reconstruct a significant number of connections within the mushroom body, with denser predictions demonstrating a higher fidelity to the original network structure.

Furthermore, we present the visualization of the embeddings of KCs by tSNE in Figure 7C, showing three clear main clusters (purple: *α*^*I*^*/β*^*I*^ neurons; red: *γ* neurons; blue:*α/β* neurons). More importantly, our model not only successfully identifies clusters within these KCs, but also reveals the presence of multiple sub-clusters within each cluster. This observation underscores the capability of our model to discern sub-types that exhibit distinct properties, thus providing valuable insights into the inherent heterogeneity within the neuronal population.

Hierarchical clustering (Figure 7D) of KCs using scores of Figure 7B(3) therefore resolves KCs into three main types. Furthermore, this approach successfully identifies the previously reported subtypes and even reveals isolated subtypes that have not yet been characterized. These compelling observations provide strong support for the assertion that our model serves as a valuable tool for differentiating functional types of neurons through its ability to learn and leverage topological features within brain circuit networks.

## Conclusions

In this paper, we introduce Graph Neural Networks (GNNs) to the neuron-level connectome, thereby establishing a novel model for inferring neuronal circuit connections. Our proposed method, called NCPNet, offers a simple yet effective approach for learning graph-topological features. From experiments, we can draw the following conclusions:

- Our model presents outstanding connection prediction capability even with small established connections. This point is important because identifying synaptic connections costs much resource [Dorkenwald et al., 2022].
- Our model can learn meaningful low-dimensional representations for a brain circuit network, which can be used for neuron clustering analysis.
- Our model can identify the functional sub-types of neurons only using connection data.

We believe that the proposed model has a good potentiality to improve the efficiency of the synaptic partner assignment procedure, and potentially promotes larger brain connectome reconstruction in the future.

## Acknowledgements

We are very grateful for the assistance provided by Prof. Anan Li from Huazhong University of Science and Technology and Dr. Jingpeng Wu from Princeton University in this work. This work is partially supported by the National Natural Science Foundation of China (62206202, 62225113), China Posdoctoral Science Foundation (2022M712461), Artificial Intelligence Innovation Project of Wuhan Science and Technology Bureau (No. 2022010702040070), The Fundamental Research Funds for the Central Universities(2042022kf1043)

## Supplementary Information

### 1 Datasets

We preprocessed the original data from Cook et al. [2019] and Scheffer et al. [2020] and converted them into an easily accessible and adaptable format by PyTorch Geometric data loaders [Fey and Lenssen, 2019]) for encouraging further study in the graph learning and neuroscience community. The detailed implementation please see the site ^1^.

**Figure S1:**
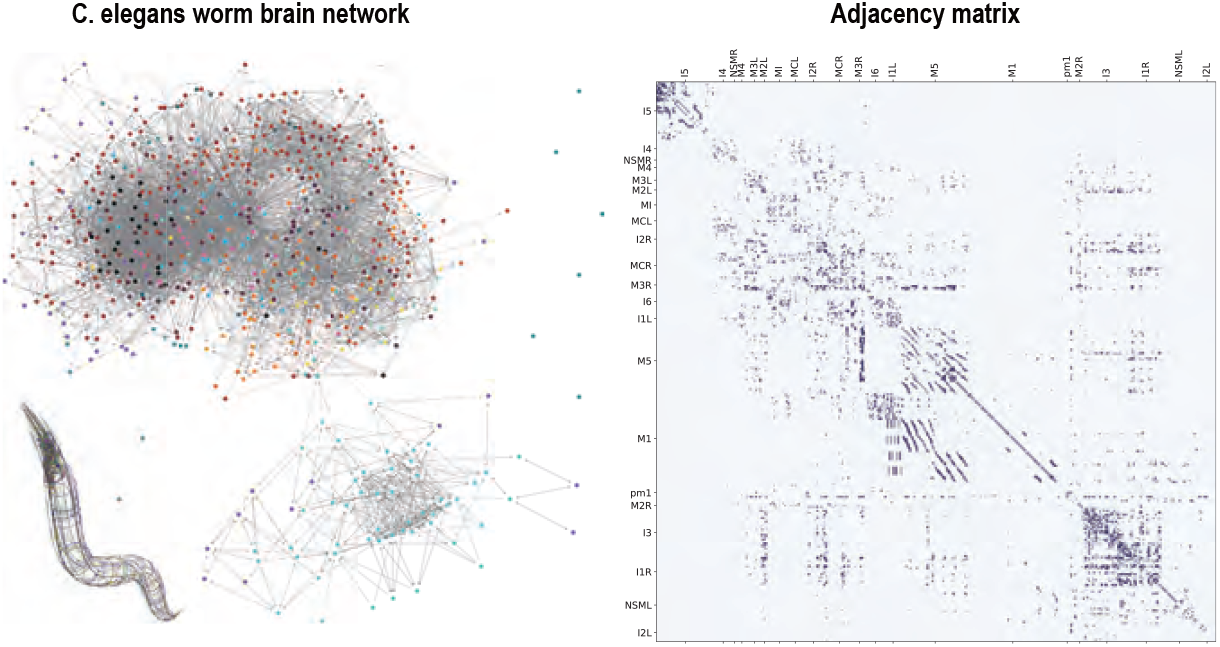
(Left) The visualization of the *C. elegans* connectome. The connectome stems from a male *C. elegans* worm. We use spring graph layout algorithm to draw this figure. (Right) The adjacency matrix of 598 neurons.

**Figure S2:**
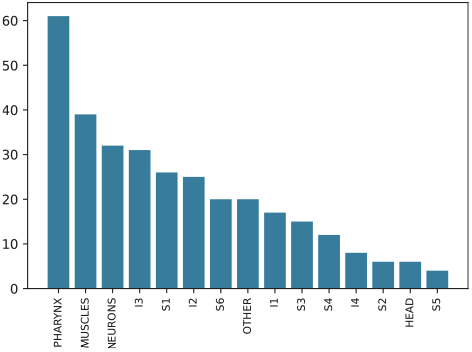
The statistic of neuron types of the *C. elegans* connectome Cook et al. [2019].

**Figure S3:**
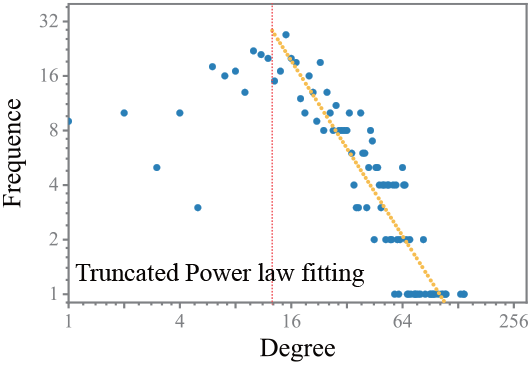
Degree distribution of the *C. elegans* connectome.

#### Type information

A type represents a specific function or region of a neuron. For example, ‘MUSCLE’ in Figure S2 denote a muscle cell for movement. the *C. elegans* connectome contains 19 types. *Drosophila* hemibrain connectome is more complex. Its neuron types are identified by multiple methods, including lineage based on cell body fibers Ito et al. [2014], computational model Costa et al. [2016].

**Figure S4:**
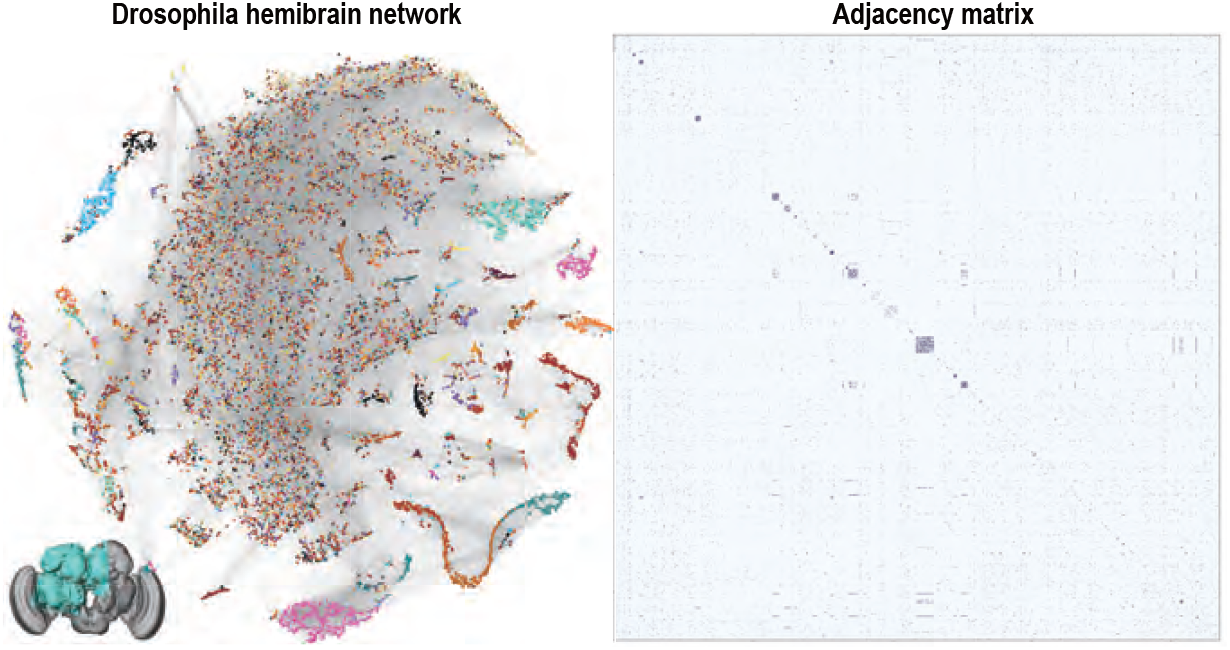
(Left) The visualization of the *Drosophila* hemibrain connectome. The connectome stems from the half brain of a fruit fly. We use NCPNet+tSNE as an efficiency graph layout algorithm to draw this figure. (Right) The adjacency matrix of 21793 neurons.

**Figure S5:**
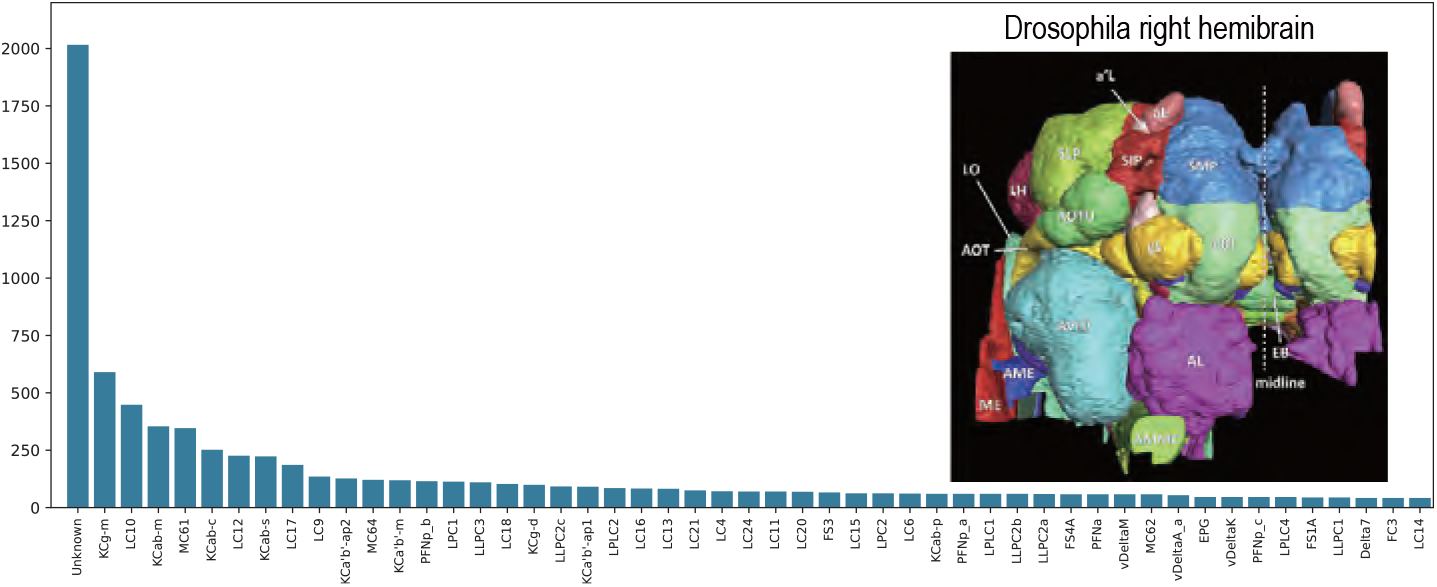
The statistic of top-50 neuron types of the *Drosophila* connectome.

**Figure S6:**
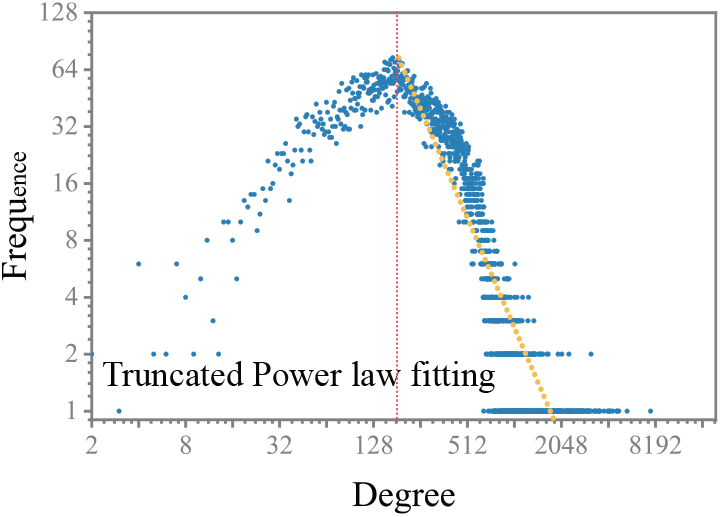
Degree distribution of the *Drosophila* connectome.

### 2 Hyperparameter settings

We use YAML configuration to control the hyperparameter in our implementation. Then we grid-searched all hyperparameters and the optimal settings are shown in Table S1. Note that we exclude AA and CN because they are hyperparameter-free methods.

**Table S1:**
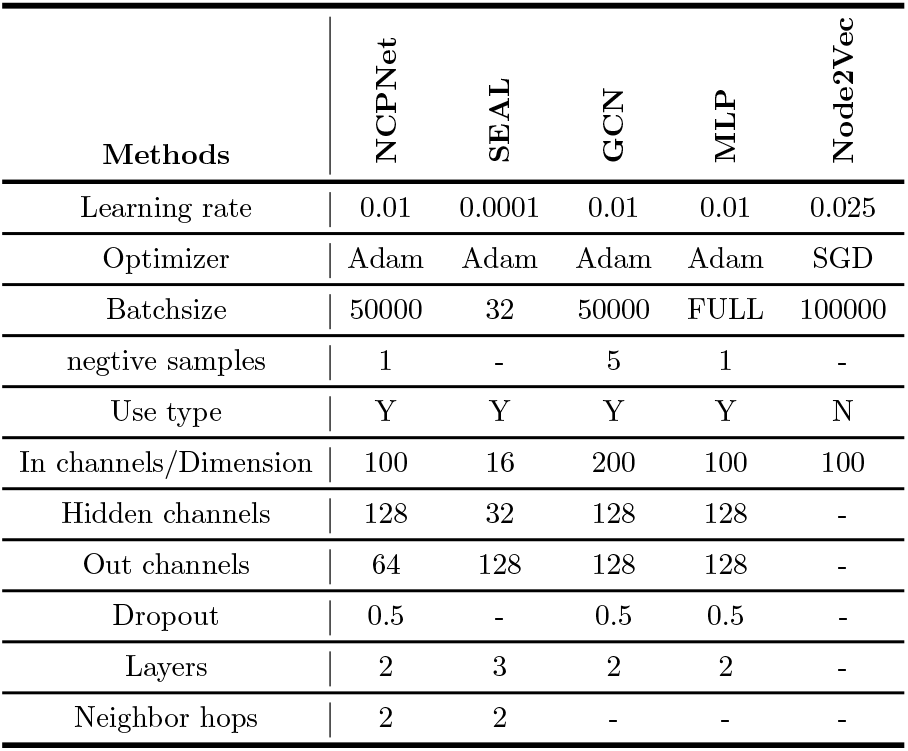
Hyperparamter settings in our experiments.

| Methods | NCPNet | SEAL | GCN | MLP | Node2Vec |
| --- | --- | --- | --- | --- | --- |
| Learning rate | 0.01 | 0.0001 | 0.01 | 0.01 | 0.025 |
| Optimizer | Adam | Adam | Adam | Adam | SGD |
| Batchsize | 50000 | 32 | 50000 | FULL | 100000 |
| negative samples | 1 | - | 5 | 1 | - |
| Use type | Y | Y | Y | Y | N |
| In channels/Dimension | 100 | 16 | 200 | 100 | 100 |
| Hidden channels | 128 | 32 | 128 | 128 | - |
| Out channels | 64 | 128 | 128 | 128 | - |
| Dropout | 0.5 | - | 0.5 | 0.5 | - |
| Layers | 2 | 3 | 2 | 2 | - |
| Neighbor hops | 2 | 2 | - | - | - |

### 3 Experiments

#### 3.1 Connection prediction

We present the accuracy, recall and precision performance similar to Figure 3A. We can observe that NCPNet exhibits good performance on accuracy and recall too. The performance on precision is lower than heuristic methods when use less training connections. The higher precision of the heuristic approach is attributed to its prediction of node connectivity by calculating the number of shared topological connections between nodes. This inherent feature allows for selectivity in predicting nodes with multi-hop connectivity, leading to a higher precision rate. However, if the connectivity between nodes is poor, the number of shared topological connections will be zero, resulting in a loss of predictive capability.

**Figure S7:**
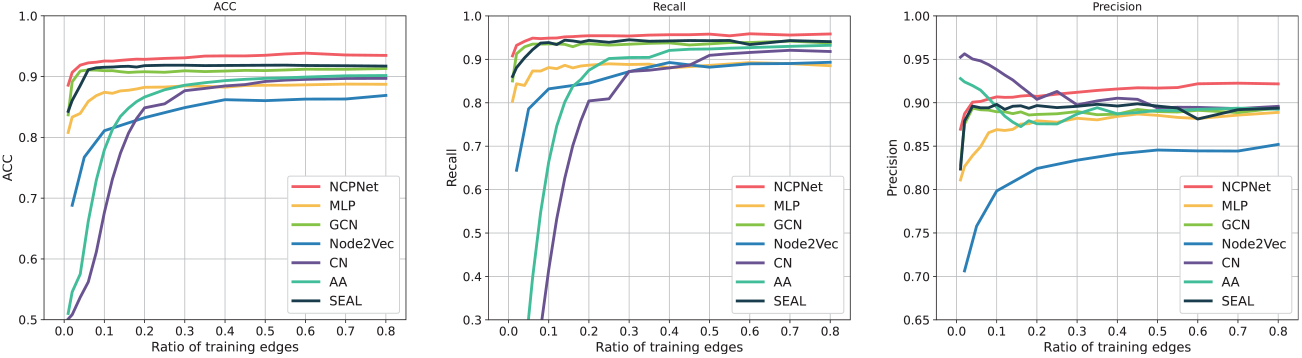
Additional numerical results:1)Accuracy, 2)Recall, 3)Precision versus training data ratio on *Drosophila* hemibrain network.

**Figure S8:**
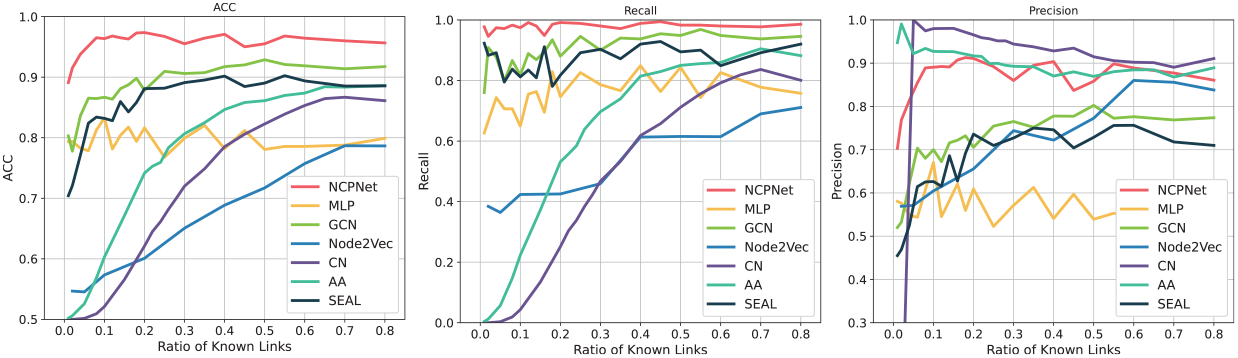
Additional numerical results:1)Accuracy, 2)Precision, 3)Recall versus training data ratio on *C. elegans* worm brain network.

#### 3.2 More prediction score analysis

Similar to Figure 6, we conducted additional predicting cases on different neurons. Due to the lack of neuronal skeleton information in the *C. elegans* worm brain network, we focused our attention more on *Drosophila* hemibrain network. We provide pre-trained model at our code repository ^2^.

**Figure S9:**
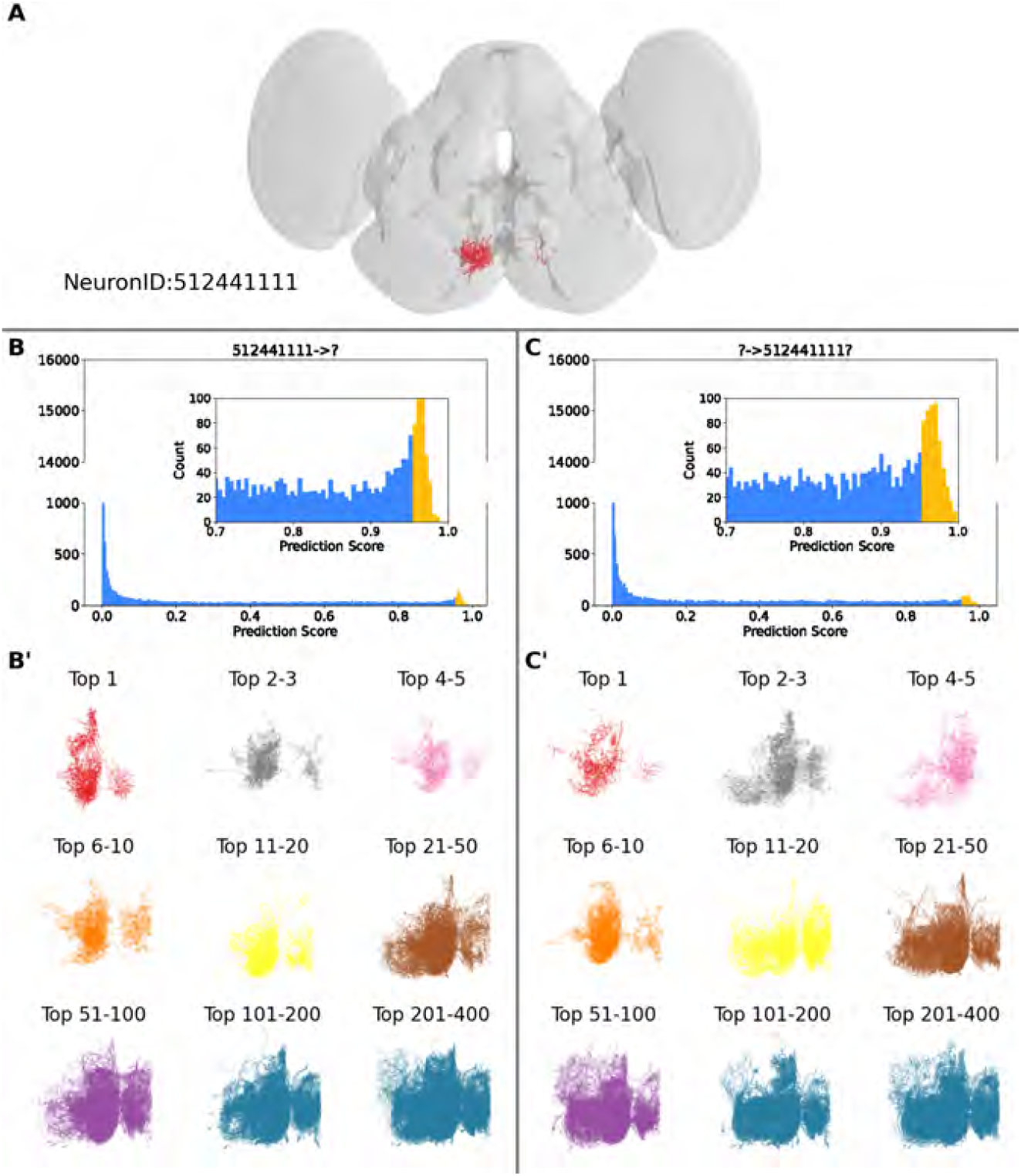
NCPNet prediction scores on *Drosophila* connectome using 5% connections. A: Neuron 512441111 in the fly brain volume. B: The raw score distribution of predicting (*v*_512441111_, ?), in which 512441111 is the pre-synaptic neuron. B’: The 9 groups of the top 400 hits of B. C: The raw score distribution of predicting (?, *v*_512441111_), in which *v*_512441111_ is the post-synaptic neuron. C’: The 9 groups of the top 400 hits of C.

**Figure S10:**
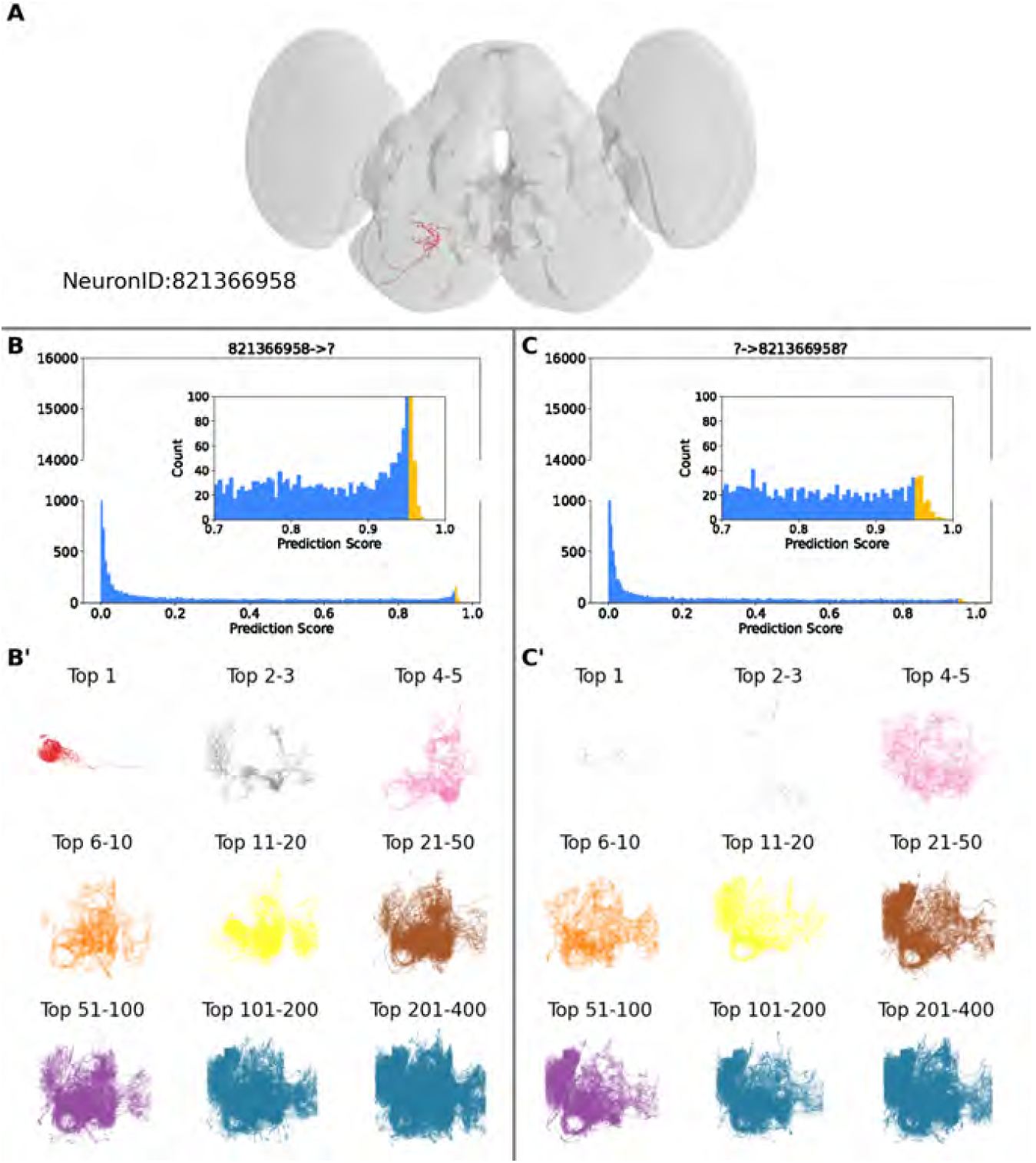
NCPNet prediction scores on *Drosophila* connectome using 5% connections. A: Neuron 821366958 in the fly brain volume. B: The raw score distribution of predicting (*v*_821366958_, ?), in which 821366958 is the pre-synaptic neuron. B’: The 9 groups of the top 400 hits of B. C: The raw score distribution of predicting (?, *v*_821366958_), in which *v*_821366958_ is the post-synaptic neuron. C’: The 9 groups of the top 400 hits of C.

**Figure S11:**
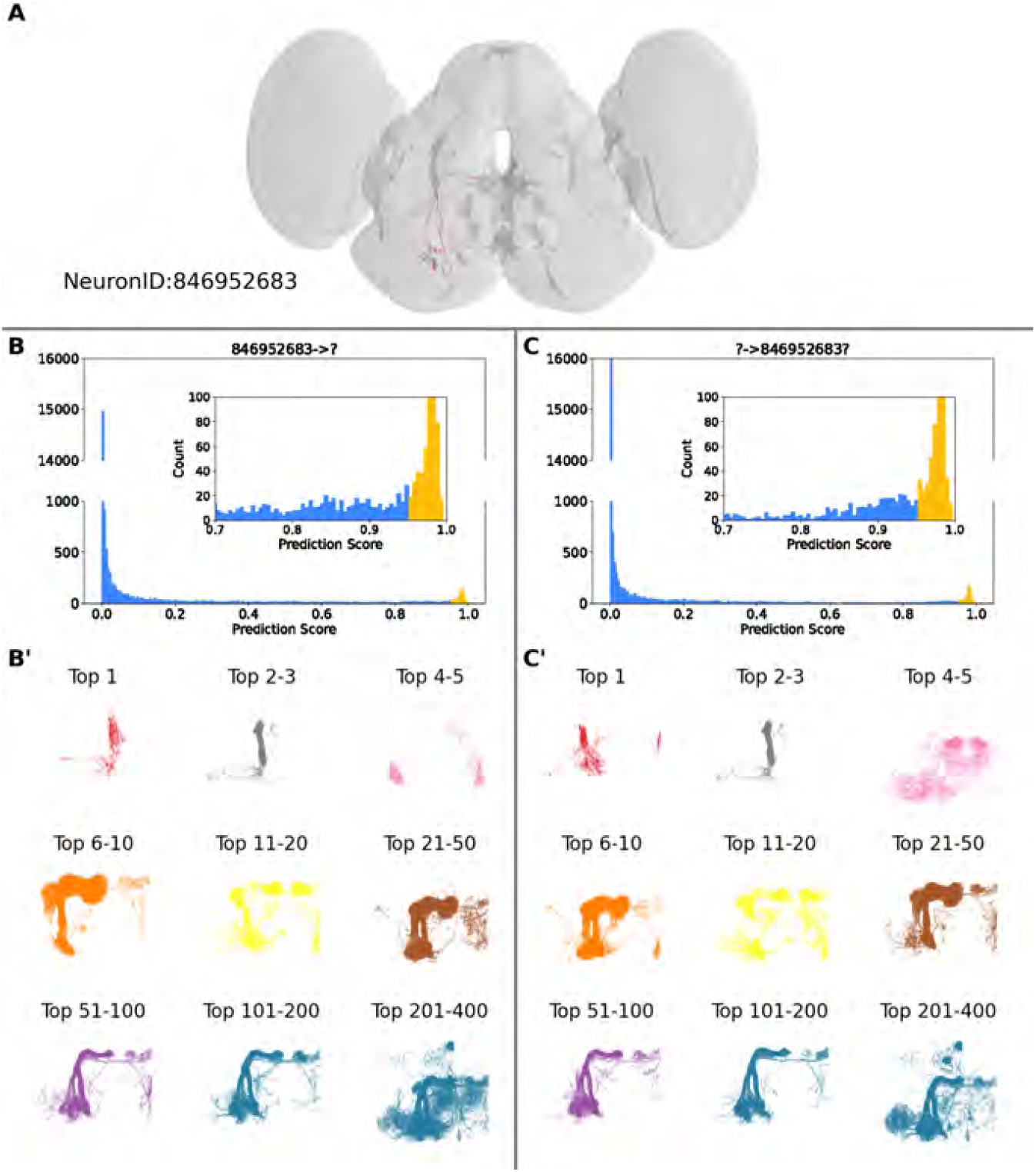
NCPNet prediction scores on *Drosophila* connectome using 5% connections. A: Neuron 846952683 in the fly brain volume. B: The raw score distribution of predicting (*v*_846952683_, ?), in which 846952683 is the pre-synaptic neuron. B’: The 9 groups of the top 400 hits of B. C: The raw score distribution of predicting (?, *v*_846952683_), in which *v*_846952683_ is the post-synaptic neuron. C’: The 9 groups of the top 400 hits of C.

**Figure S12:**
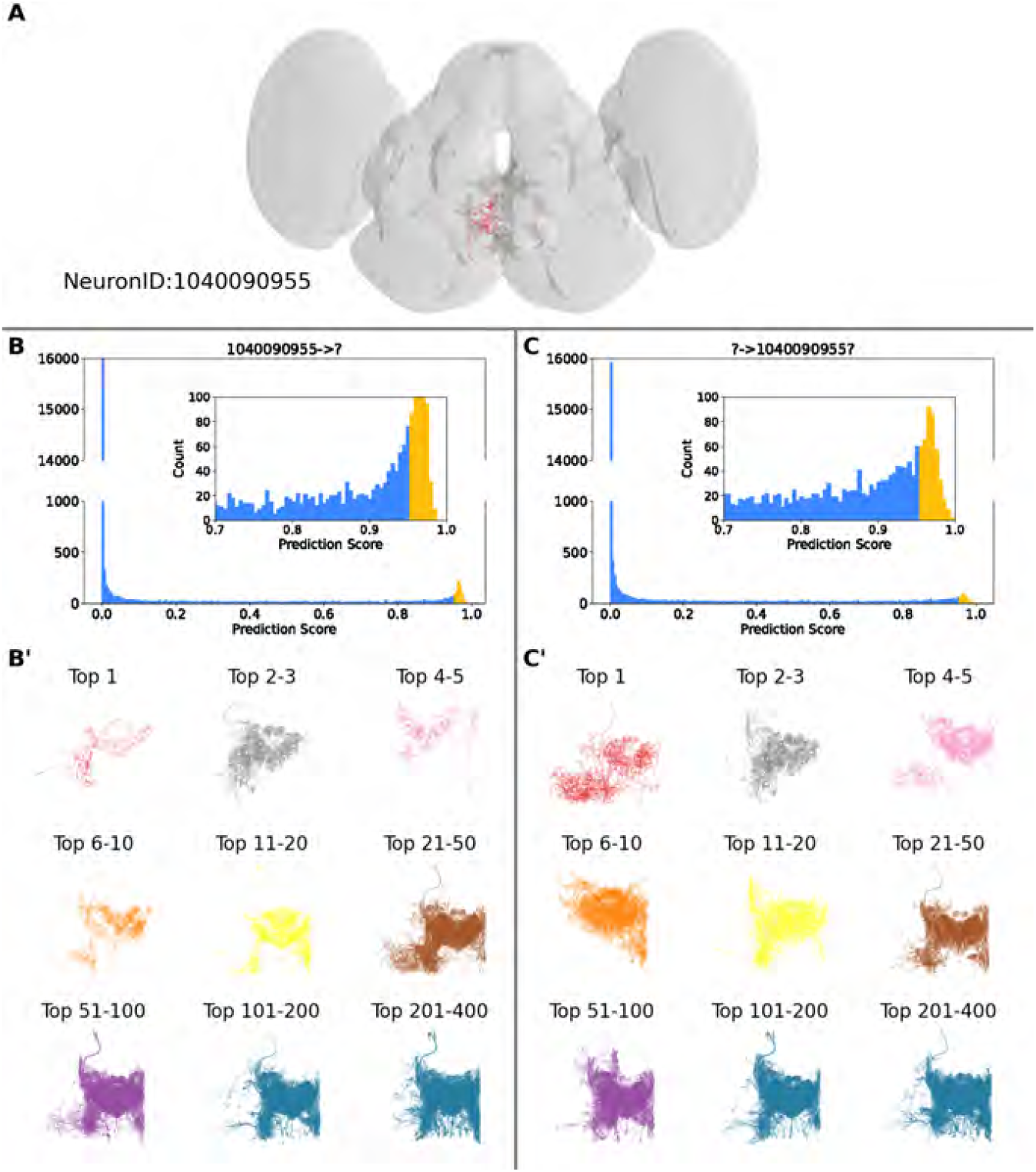
NCPNet prediction scores on *Drosophila* connectome using 5% connections. A: Neuron 1040090955 in the fly brain volume. B: The raw score distribution of predicting (*v*_1040090955_, ?), in which 1040090955 is the pre-synaptic neuron. B’: The 9 groups of the top 400 hits of B. C: The raw score distribution of predicting (?, *v*_1040090955_), in which *v*_1040090955_ is the post-synaptic neuron. C’: The 9 groups of the top 400 hits of C.

## Footnotes

1 https://www.janelia.org/project-team/flyem/hemibrain

2 https://github.com/mxz12119/NCPNet

3 https://wormwiring.org/

4 https://github.com/DeepGraphLearning/graphvite

5 https://networkx.org/

6 https://neuprint.janelia.org/

1 https://github.com/mxz12119/NCPNet

2 https://github.com/mxz12119/NCPNet/tree/main/examples/Predictingcase.ipynb

